# A comprehensive and scalable database search system for metaproteomics

**DOI:** 10.1101/053975

**Authors:** Sandip Chatterjee, Gregory S. Stupp, Sung Kyu (Robin) Park, Jean-Christophe Ducom, John R. Yates, Andrew I. Su, Dennis W. Wolan

## Abstract

**Background:** Mass spectrometry-based shotgun proteomics experiments rely on accurate matching of experimental spectra against a database of protein sequences. Existing computational analysis methods are limited in the size of their sequence databases, which severely restricts the proteomic sequencing depth and functional analysis of highly complex samples. The growing amount of public high-throughput sequencing data will only exacerbate this problem. We designed a broadly applicable metaproteomic analysis method (ComPIL) that addresses protein database size limitations.

**Results:** Our approach to overcome this significant limitation in metaproteomics was to design a scalable set of sequence databases assembled for optimal library querying speeds. ComPIL was integrated with a modified version of the search engine ProLuCID (termed “Blazmass”) to permit rapid matching of experimental spectra. Proof-of-principle analysis of human HEK293 lysate with a ComPIL database derived from high-quality genomic libraries was able to detect nearly all of the same peptides as a search with a human database (~500x fewer peptides in the database), with a small reduction in sensitivity. We were also able to detect proteins from the adenovirus used to immortalize these cells. We applied our method to a set of healthy human gut microbiome proteomic samples and showed a substantial increase in the number of identified peptides and proteins compared to previous metaproteomic analyses, while retaining a high degree of protein identification accuracy, and allowing for a more in-depth characterization of the functional landscape of the samples.

**Conclusions:** The combination of ComPIL with Blazmass allows proteomic searches to be performed with database sizes much larger than previously possible. These large database searches can be applied to complex meta-samples with unknown composition or proteomic samples where unexpected proteins may be identified. The protein database, proteomics search engine, and the proteomic data files for the 5 microbiome samples characterized and discussed herein are open source and available for use and additional analysis.

## Background

Precise characterization of the protein components of a cell or tissue is a powerful technique for assessing a functional state of a system, as direct detection of expressed gene products can enable more accurate systems-level analysis of a cellular state than the information gathered from sequencing genes or transcripts alone [1, 2]. The method of choice for modern, discovery-oriented proteomics experiments is tandem mass spectrometry (MS/MS), which relies on peptide mass, charge state, and fragmentation to identify proteins in a sample.

MS/MS data is commonly analyzed and assigned to peptide sequences using software such as SEQUEST, Mascot, and ProteinProspector [3–5]. A critical component in applying these proteomic data analysis tools is the choice of an adequate collection of protein sequences. Peptide candidates for each mass spectrum are selected from this protein sequence database and scored against experimental MS/MS data. In this approach, high-scoring peptide candidate matches are chosen as peptide identifications for spectra after rigorous statistical filtering and post-processing [6–8].

For each of these analysis tools, a peptide sequence must be present in the chosen sequence database for it to be identified in a biological sample. Spectra corresponding to peptides that are not present in the sequence database – including slight variants of database peptides – will be left unidentified or incorrectly identified. The use of an inappropriate or incomplete protein sequence database for a sample can result in erroneous conclusions following protein identification. For instance, a recent study reanalyzed an existing *Apis mellifera* (honey bee) proteomic dataset that purportedly contained viral and fungal proteins that were implicated in honey bee colony collapse. Upon reanalysis using a larger, more inclusive proteomic search database, the spectra corresponding to viral and fungal peptides were reassigned to better-scoring honey bee peptides, calling into question the original authors’ findings [9].

When a heterogeneous biological sample, such as a complex environmental or microbiome sample, is assessed by current proteomic analysis methods there is the potential for many mass spectra to remain unassigned and subsequently for many proteins from the sample to be left unidentified. Unfortunately, current proteomic analysis algorithms can not perform extremely large database searches in a practical time frame, restricting searches to more limited protein search databases. Recent high-throughput genomic sequencing efforts have begun to characterize variation at the gene and protein sequence levels, leading to a deluge of publicly available protein sequence data [10–15]. The breadth of these types of studies and the data generated are potentially valuable for thorough proteomic data analysis [2, 16]. However, such extensive protein sequence databases derived from enormous panels of organisms with many protein variants will easily overwhelm currently available proteomic search methods that were historically designed to interrogate single proteomes. An accessible computational search method incorporating a comprehensive, rapid, and scalable protein sequence database comprised of all known public protein sequences would greatly advance the burgeoning field of metaproteomics, allowing for proteomic characterization of highly complex, unculturable biological samples, or for any biological system where significant protein sequence variation may be present.

To address these limitations, we designed a method that utilizes high-performance peptide and protein databases, scalable to an essentially unlimited number of protein sequences, “ComPIL” (Comprehensive Protein Identification Library), which is tightly integrated with a modified version of ProLuCID [17], “Blazmass”. Additionally, we compiled a ComPIL database consisting of an enormous amount of high-quality genomic sequence information (approximately 500x the size of the human proteome). Here, we validate and evaluate the sensitivity and specificity of ComPIL using human and individual bacterial samples, reanalyze public proteomics data for discovery of new biomedical knowledge, and then demonstrate the method’s utility in improving protein and functional annotation on a number of human microbiome samples.

## Results

### Organizing large amounts of protein information into optimized search databases

To account for the immense size of a comprehensive protein database and optimize for the retrieval of information needed for efficient peptide-spectrum scoring, we organized our protein data into three distributed NoSQL databases (**Figure 1A**) implemented using MongoDB (https://www.mongodb.org/). Protein sequences were stored in a database termed ProtDB and digested *in silico* using trypsin specificity. The resulting peptide sequences were grouped by identical peptide mass or sequence and stored in databases MassDB or SeqDB, respectively (**Figure S1**). This design is readily scalable as more protein sequence information becomes available (see **Methods** for details on database organization).

**Figure 1.**
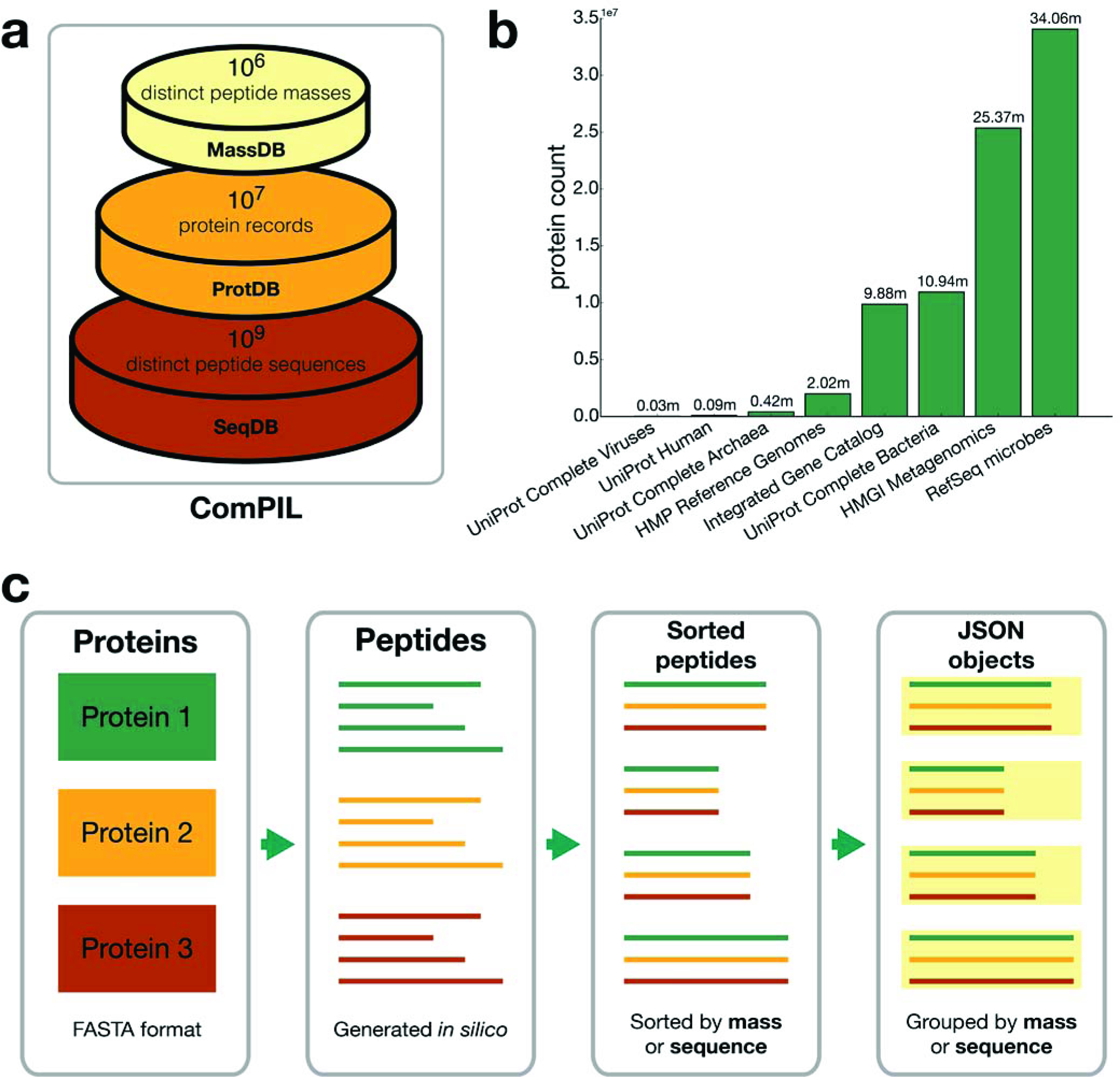
Design, components, and generation of ComPIL databases. **(a)** ComPIL utilizes 3 databases that are generated from an input protein FASTA file. MassDB contains peptide sequences organized by distinct mass: ProtDB contains protein information; SeqDB contains distinct peptide sequences along with their parent proteins (mapped to ProtDB). **(b)** Public protein repositories and numbers of proteins incorporated into ComPIL. Numbers shown above columns are in millions, **(c)** 1) Protein data from various repositories (shown in b) were grouped together in FASTA format. Protein records were imported into ProtDB. 2) Proteins were *in silico* digested to peptides using trypsin specificity. 3) Peptides were sorted by sequence or by mass to group peptides with identical sequences or masses together, respectively. 4) Peptides with identical sequences or masses were grouped into JSON objects which were imported into MongoDB as SeqDB or MassDB. respectively. See **Methods** for implementation details.

Owing to the recent interest in microbiome studies and the tremendous improvements in sequencing technologies, a wealth of microbial genome data is now available from public sequence repositories. A ComPIL database was generated by amassing protein sequence data from a number of large, public sequencing projects (**Figure 1B**, **Methods**, and **Table S1**). In order to assess false discovery rate (FDR) using the target-decoy approach, protein sequences were reversed and concatenated with their original protein records to produce a large FASTA file (available for download, see **Methods**). This data was organized into ProtDB, MassDB, and SeqDB using an efficient data processing pipeline, producing a total of 165.6 million protein records and roughly 4 billion unique tryptic peptide sequences (**Figure 1C** and **Methods**). Only 0.7% of all peptide sequences mapped to both real and decoy proteins (**Figure S2**), which is unlikely to affect the filtering of proteins due to the filtering requirement of matching 2 peptides per protein. The vast majority of peptide sequences (84%) map to three or fewer parent protein sequences (**Figure S3**); therefore, each individual peptide sequence in the database represents a substantial amount of new protein information. However, despite the increased search space provided by ComPIL, the library does not approach all possible peptide amino acid sequences required for *de novo* sequencing, which is many orders of magnitude larger.

### Database searching and scoring

Modern proteomic search software typically incorporate a pre-processing step on the input FASTA protein database file in order to speed up searches. For example, ProLuCID pre-processes the FASTA file by *in silico* digesting all proteins, grouping the peptides by mass, and storing them in a local SQLite database, while Crux similarly utilizes tide-index, storing the data in a binary file on disk [17, 18]. These greatly speed up searches for small to medium sized databases; however, they do not scale to handle the billions of distinct peptide sequences in a ComPIL search.

In this work, data was stored using MongoDB, which provides the primary benefits of speed and scalability over existing pre-processing techniques. MongoDB (and other NoSQL-type databases) allow simple “horizontal” scaling to a computational cluster of machines, meaning that a database can grow indefinitely by adding more machines while the querying and retrieval workload are distributed across all machines. This design is optimal for distributed analysis as an MS dataset can be divided into chunks that are processed independently on cluster nodes, with individual nodes each querying the same proteomic database over a local network. Using this method, a local copy of the proteomic search database does not have to be generated or copied onto each node of a cluster. ComPIL was deployed as a sharded database across 8 networked Linux servers (see **Methods** for implementation details).

We modified the ProLuCID query engine to allow integration with ComPIL databases, or any other MongoDB databases. This was implemented using the Java MongoDB driver, a high-level driver that allows asynchronous database operations. Individual search threads are parallelized within one process, all utilizing the same MongoDB connection. As is typical for many proteomics database search engines, a calculated mass from a precursor ion is first used to query for a list of possible peptide candidates. The peptides are returned as a stream, implemented using Java's Iterable interface, and the data is subsequently streamed from the database to the worker, reducing memory usage on the worker.

A variety of scores are calculated for each peptide candidate-spectrum pair in order to determine the best possible match. Blazmass calculates XCorr, a cross-correlation score ranging from 0 to 10, for each peptide candidate-spectrum pair and ranks peptide candidates for each spectrum in descending order by XCorr value. This peptide candidate ranking allows a second score (DeltaCN), a normalized score ranging from 0 to 1 representing the XCorr difference between the first and second best peptide matches, to be calculated for each spectrum. Additionally, a Z-score is calculated using the XCorr of the top hit, which aims to measure if the top hit is significantly different from the other candidates [17]. A direct correlation between an increase in deviation from top score to other possible hits in the list increases the likelihood and confidence that the peptide is a true hit. A combination of these scores is typically used to assess the quality of a peptide-spectrum match using DTASelect2, or other postprocessing software such as Percolator [7, 8].

The integration of Blazmass with ComPIL allows for efficient proteomic scoring against extremely large search spaces. We attempted to benchmark the speed and accuracy of this software against SearchGUI, a software package allowing for the simultaneous use of 8 different proteomic search engines [19]. Despite extensive effort, we were only able to search a small section of a mass spectrometry file with the ComPIL database using one of the proteomic search engines: Comet [20]. The primary reasons for failure of the other search engines stemmed from insufficient RAM required for the database indexing step or significant limitations in search speeds that prevented timely results. We compared the performance of Comet to Blazmass on 100 scans, resulted in a filtered PSM agreement of 100%, as defined by both search engines matching the same peptide without post-translational modifications as the top match, however, Comet was ~150x slower (see **Methods** for details).

### Human peptides are correctly identified within the expanded search space of ComPIL

A significant concern with larger protein databases is the accuracy of peptide-spectrum matches [16, 21, 22]. In the ComPIL database, the overall mass distribution of peptides is similar to peptides derived from the human proteome, but the large database contains ~500x as many peptides for any given calculated peptide mass, resulting in a large number of potentially incorrect matches for each mass spectrum. To assess the accuracy of a ComPIL search, we designed an experiment in which tandem MS data from a known proteome would be searched against both ComPIL and a standard, single-organism protein database (**Figure 2A**). The peptide-spectrum matches (PSMs) from both searches were compared to assess agreement in peptide sequence assignment.

**Figure 2.**
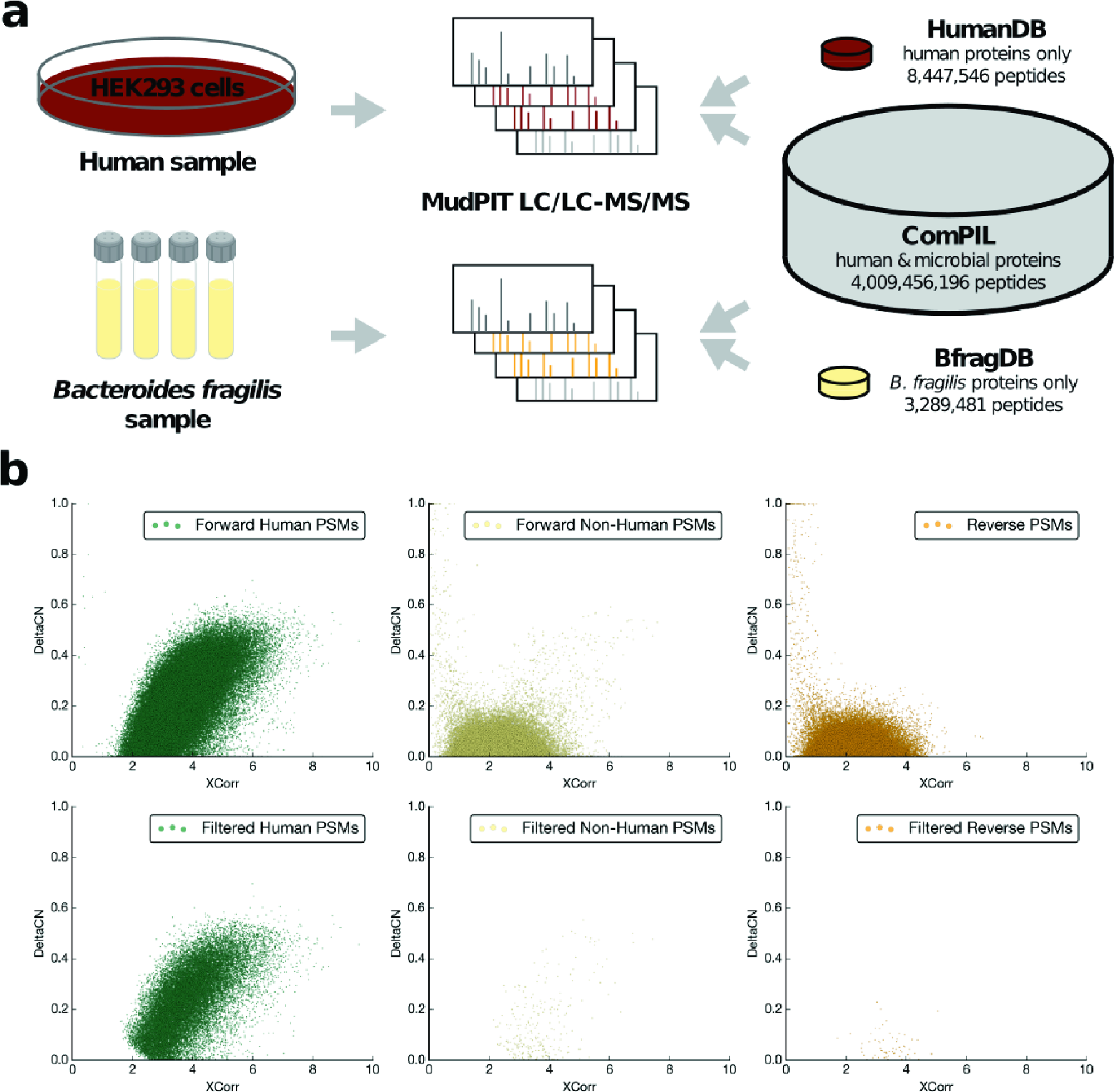
Validation of ComPIL with Blazmass searches using human and *B. fragilis* samples. **(a)** Human and *B. fragilis* proteins were extracted and tandem MS data was collected using MudPIT. The datasets were searched against ComPIL and either the human proteome or the *B. fragilis* proteome. **(b)** ComPIL searches of HEK293 cells. Unfiltered PSMs (top) and filtered PSMs (after filtering at 1% FDR with DTASelect2. bottom) are shown categorized as forward human matches (left), forward non-human matches (middle) or reverse (*e.g.*, decoy) protein matches (right).

We collected tandem MS data from a human HEK293 cell lysate using MudPIT [23] and searched the resulting dataset against both the human proteome alone and the ComPIL database, which includes the human proteome as a small subset (0.11% of ComPIL proteins). Plots of Blazmass-assigned DeltaCN vs. XCorr values for PSMs from the ComPIL search revealed that spectra matching “human” peptides clustered distinctly from both “non-human” (*i.e.*, generally presumed incorrect for a human sample) and reverse “decoy” (*i.e.*, false positive) peptide matches (**Figure 2B**). Erroneous non-human and reverse PSMs clustered together, with both groups having similarly low median XCorr (2.24 non-human, 2.22 reverse) and DeltaCN (0.03 non-human, 0.03 reverse) values. The clustering of correctly and incorrectly identified PSM data closely matches that observed for “forward” and “reverse” PSMs in a target-decoy search against the human proteome (**Figure S4**), illustrating that the ComPIL search of a human sample is correctly differentiating human and non-human PSMs.

We used DTASelect2 to filter PSM data based on a maximum 1% protein FDR and minimum 2 peptide identifications per protein. The FDR needs to be carefully controlled at the protein level, as often the protein FDR can exceed the PSM FDR as the database size increases [21]. Blazmass scored each HEK293 peptide spectrum against approximately 500x as many peptide candidates in the ComPIL search as in the human proteome search, yet the accuracy of high-quality, filtered peptide matches was strikingly high in the ComPIL search. We were able to identify the same peptide for 99.7% of 25,627 filtered spectra appearing in both filtered search result datasets. Surprisingly, using the expanded search space afforded by the ComPIL database, we also identified peptides corresponding to human adenovirus 5 proteins E1A and E1B (**Table S2**), which are known contaminants and historical artifacts of transformation of the HEK293 cell line [24]. This finding demonstrates the power of ComPIL in elucidating important, unexpected protein information that current proteomic search methods and narrow-spectrum databases will be incapable of identifying.

While PSM accuracy remained high, we observed a 15% decrease in sensitivity (number of PSMs appearing in filtered data for human proteome vs. ComPIL search), as has previously been predicted for large database searches [22]. This loss in sensitivity can be explained by a more stringent filtering of XCorr scores in the ComPIL search (median = 3.74) compared to the human proteome search (median = 3.51), as well as a decrease in deltaCN (median = 0.236) compared to the human proteome search (median = 0.440) (**Figures 2B** and **S4**). An overlay of score distributions (XCorr vs. DeltaCN) for peptides found in the ComPIL and human proteomes (**Figure S5**) implicates an overall decrease of DeltaCN values in the loss in peptide identification sensitivity. This decrease, expected as database size grows, makes differentiation of true and false peptide matches more difficult. Notably, 531 additional peptides (a 2.6% increase) were found only in the ComPIL search and can be attributed to several possible sources, such as false hits, proteins from non-human species sharing high sequence similarity with humans, incorrectly annotated proteins, and/or contaminants such as HAdV5 (**Figure S6**).

### Bacterial peptides are accurately identified within the large search space of ComPIL

We next investigated whether human peptides might somehow be easier to identify than microbial peptides, as microbial peptides constitute the vast majority of ComPIL and share limited sequence identity with human peptides (**Figure 2A**). As a second test of PSM accuracy, we performed a similar experiment by collecting multidimensional protein identification technology (MudPIT) data from a *Bacteroides fragilis* cell lysate and searched the data against both a *B. fragilis* only protein database and ComPIL. The *B. fragilis* proteomic database, like the human database, constitutes a very small portion of ComPIL (representing 0.08% of all ComPIL peptides). As in the human proteomics experiment, we found that the vast majority of 14,812 PSMs (98%) appearing in both filtered datasets mapped to the exact same peptide sequence, again highlighting the accuracy of ComPIL-Blazmass as compared to a database search of a more restricted proteome. We again observed a decrease in sensitivity with the larger search space, with a 20% reduction in the number of filtered PSMs appearing in the final output.

### Detection of low-abundant pathogen proteins in human proteomic samples with ComPIL

Due to the unexpected detection of adenovirus 5 proteins within the HEK293 cellular lysates, we assessed if unbiased ComPIL searches would be capable of identifying low abundance pathogenic proteins from a sample of human cells infected with Influenza A. Specifically, we obtained publicly available proteomic data collected from an experiment in which Calu-3 human lung cancer cells were infected with a wild-type Influenza A strain and harvested after 0, 3, 7, 12, 18, and 24 hours for MS-based proteomic data collection (ProteomeXchange: PXD002385). Importantly, samples for the proteomics data collection were not enriched for Influenza proteins and thus unbiased towards the detection of peptides corresponding to Influenza. ComPIL searches detected Influenza A virus peptides in the datasets beginning at 7 h post-infection and represented the second most abundant genus of proteins in the 18 h sample after human with approximately 5% of the total spectral counts (**Figure 3**). In summary, we observe a direct relationship between an increase in infection time of the Calu-3 cells with an increase in number and spectral counts of the Influenza A peptides.

**Figure 3.**
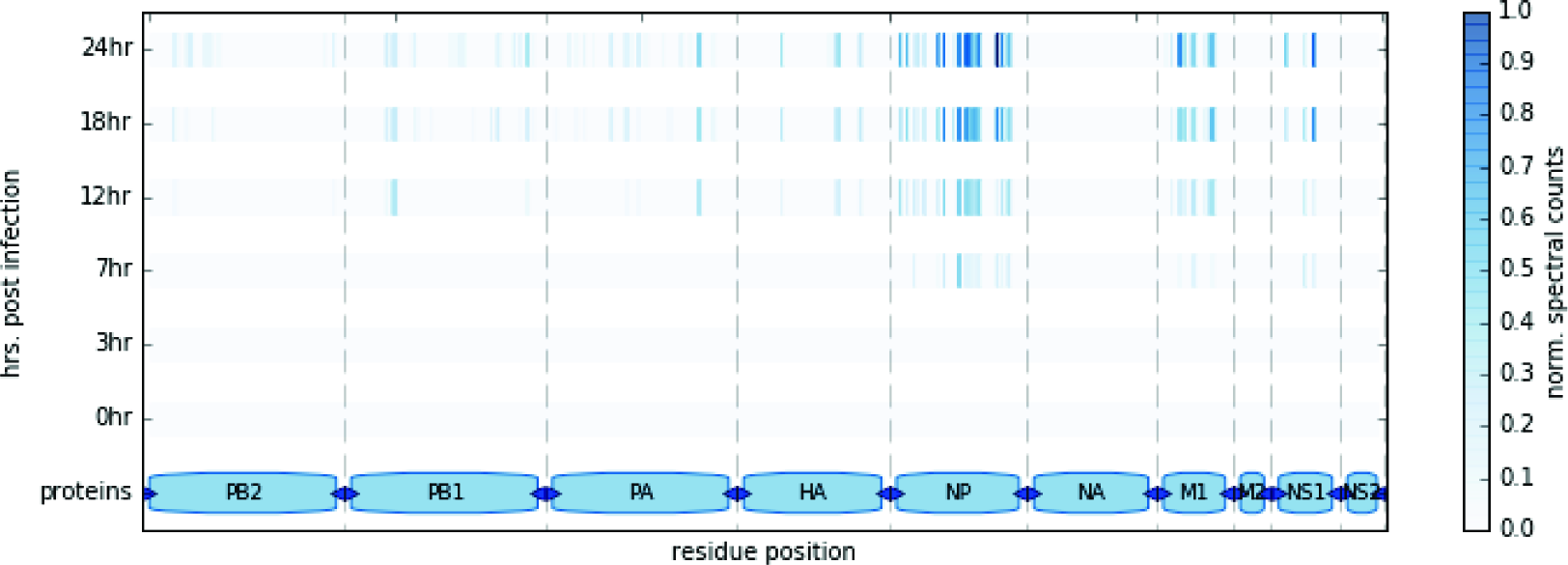
Post-infection detection of Influenza A peptides in Calu-3 cells searched using the ComPIL database. Detected Influenza peptides are shown mapped to their location within the Influenza A/Anhui/1/2013 proteome as a function of time. The color represents the normalized spectral counts of peptides found at each residue. Influenza A peptides could be detected from 7 h post-infection, and shown a direct relationship between infection time and relative quantitation of the peptides. A Jupyter notebook with more details about this figure is available at https://bitbucket.org/sulab/metaproteomics.

### Use of ComPIL yields significantly more peptides and protein functionalities within the human gut microbiome

We evaluated the performance and proteomic results of the ComPIL database in comparison to a limited metaproteomic search database used to characterize healthy human microbiota. A library of complete proteomes (termed “46 proteomes”) based on a previously published and validated metaproteomic sequence database [25] was generated (consisting of a small subset of ComPIL) (**Table S3**). For reference purposes, this database contains about 100x fewer peptides. MudPIT data collected from the soluble proteome fractions of four healthy human fecal samples was searched against both databases. After DTASelect filtering, we observed a significant increase in the number of PSMs identified by the ComPIL search for all four samples, when compared to the more focused “46 proteomes” database (**Figure 4A**).

**Figure 4.**
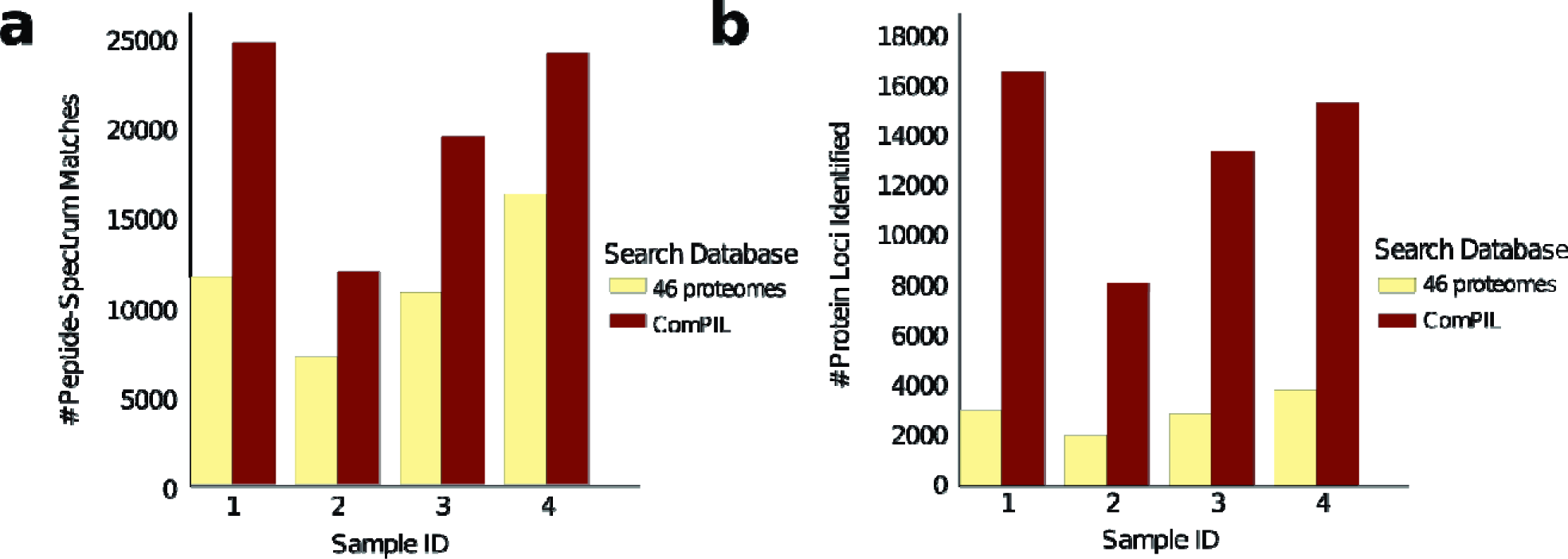
Evaluation of ComPIL-Blazmass search of a complex microbiome sample. **(a)** Filtered PSMs from four human stool proteome samples after searching each clataset with either the “46 proteomes” database or ComPIL. **(b)** Protein loci identified from four human stool proteome samples after searching each dataset with either the “46 proteomes” database or ComPIL.

Importantly, the increase in high-quality PSMs identified in our ComPIL search resulted in substantially more protein locus identifications (**Figure 4B**), using ComPIL as compared to the “46 proteomes” database search. These results are due to the increasing number of PSMs as well as to the larger number of proteins within the database the PSMs match. After filtering with DTASelect, >95% of the PSMs appearing in the smaller database search mapped to the same peptide sequence in the larger database search, showing that a limited database search is generally accurate in its peptide identification, but fails to assign a large number of spectra. An overlay of the score distributions (XCorr vs. DeltaCN) from peptides identified with ComPIL or 46 proteome searches for one fecal sample (H1_1) reveal an overall suppressed DeltaCN for the larger ComPIL database search (**Figure S7**). This results in a decrease in sensitivity, as previously shown with the proteomic searches of ComPIL against the more focused human and *B. fragilis* proteomics search databases (**Figures 3** and **S4**). However, a substantial number of MS spectra were matched to peptides only contained within the ComPIL database (**Figure S8**). Significantly, these ComPIL-only peptide matches are evenly distributed throughout the range of XCorr and DeltaCN scores and affirms the use of a large peptide database to accurately identify protein-derived peptides from a complex proteomic sample with unknown constituents in comparison to smaller focused libraries.

### ComPIL searches on human distal gut microbiome samples reveal a tremendous amount of proteomic diversity

As the ultimate goal of ComPIL is to search complex samples in an unbiased manner, we collected and prepared 5 human stool samples from healthy volunteers for identification of both intracellular and secreted microbial proteins with standard MudPIT data collection methods (see **Methods** for sample preparation details). Three technical replicates were subjected to MS MudPIT data collection and after filtering with DTASelect2 using the strict parameters of 2 peptides per protein, <10 ppm mass error, and 1% protein FDR, we observed an average of >16,000 unique peptides forming an average >9,000 protein loci per sample (**Table S4**). Importantly, our results demonstrate a vast increase in protein identifications compared to published methods on similar, related human fecal samples [25–28].

To facilitate the functional comparison of proteins detected in samples, we used CD-HIT [29] to cluster the ComPIL database at 70% sequence identity and formed “protein clusters”, which share common functionality (see **Methods** for clustering details). An average of 5,668 protein clusters was detected in each of the 15 samples with a total of 17,599 protein clusters identified across all samples (**Table S4**). Proteins from each protein cluster were annotated with GO terms using InterProScan [24]. We subsequently quantified the GO categories by the total protein cluster MS spectral counts associated with each GO term (**Figure 5**). The protein functionalities and compositions detected are similar to previous studies that employed shotgun proteomics [27, 28], in that proteins involved in energy production (GO:0055114 oxidation-reduction process), translation (GO:0003735 structural constituent of ribosome), and carbohydrate metabolism (GO:0005975 carbohydrate metabolic process) are highly abundant. However, many additional functionalities are differentially observed, such as signal-transduction (GO:0016301 kinase activity).

**Figure 5.**
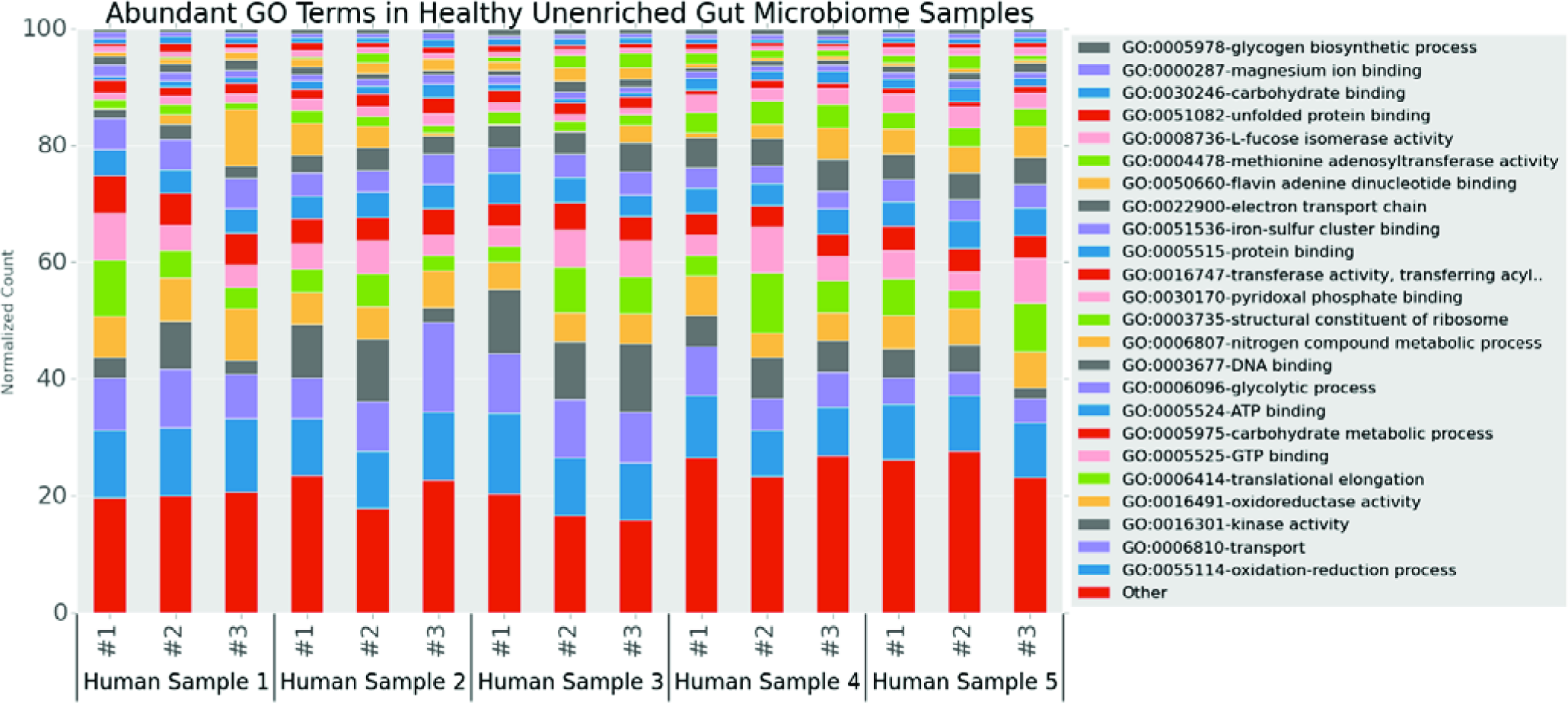
Functional annotation of five human microbiome proteome samples. Stacked bar chart showing the most abundant GO terms in each sample quantified by spectral counts. GO terms comprising the top 80% of spectral counts (on average across all samples) are shown, with the others GO terms grouped into the "Other" category. Represented are the five healthy human fecal samples subjected to metaproteomics and the three technical replicates of each sample.

### Direct identification, cloning, and exogenous expression of unannotated proteins with unknown functionalities

Many of the proteins detected in the microbiome samples have unannotated functions. Such functionalities represent potentially important proteins that will require additional investigation to determine biological relevance. To this end, we searched for proteins detected in the majority of healthy human fecal sample proteomics results with domains of unknown function (DUFs). We focused on DUF PF09861, as this protein was detected among 14 out of 15 samples and all corresponding peptides specifically mapped to a protein containing this DUF. In ComPIL, all DUF PF09861 containing proteins are flagged as “hypothetical” or “uncharacterized” and are found in the genomes of *Eubacterium* and *Clostridium* species. As a proof-of-concept downstream analysis of our unbiased, large database proteomic searches, we successfully cloned, expressed, and validated the presence of this DUF PF0986-containing protein within our healthy human fecal samples (**Figure S9** and **Methods**).

### Discussion and conclusions

The method presented here utilizing our ComPIL database integrated with the Blazmass proteomic search engine is a scalable, efficient approach for protein identification in highly complex samples potentially containing a large number of proteins. By designing the method around three databases, each with a distinct schema (**Figure S1**), each database is individually scalable and optimized for a typical workload of proteomic search queries. For example, MassDB was designed as a relatively lean, distributed database with peptide sequences organized by exact mass. We chose this design to ensure that peptide mass range-based database queries would be fast and efficient. If more proteins are added to ComPIL, new peptide sequences are appended to existing records in MassDB without creating additional database records. As a result, database search time increases linearly as ComPIL scales to accommodate additional sequences (**Figure S10**).

We therefore anticipate that computational hardware will not be a limiting factor as the ComPIL database size increases. However, we do expect a reverse correlation in peptide matching sensitivity to the increase in database size, as true PSMs will become more difficult to differentiate from false PSMs, a phenomenon which has been previously observed [22]. Notably, there are several pre-database search strategies that can be applied to address this limitation [16, 21, 30–32], such as reduction of the peptide search space with pre-filters using peptide sequence tag information or physiochemical properties (*e.g.*, isoelectric point). Similarly, post-database search strategies can assist when PSMs become difficult to differentiate, including the grouping together of redundant proteins into “meta-proteins”, which better captures the actual protein content in a sample.

Although there appears to be a loss in peptide identification sensitivity when comparing ComPIL directly against more focused, traditional database search methods, a relatively well-characterized, culturable protein sample from a HEK293 or *B. fragilis* cell culture would not typically be scored and analyzed using ComPIL. Such experiments would be analyzed using a more appropriate search database. We anticipate that ComPIL will be most beneficial for biological systems where accounting for sequence variability can quickly cause an explosion in database size and proteomic search time. This method is complementary to existing peptide sequence tag-based methods for analyzing peptide sequence variation [32, 33].

Recent proteogenomic approaches have attempted to circumvent this “big data” problem by coupling RNASeq-or metagenomic sequencing-derived protein sequence databases unique to each biological sample with subsequent proteomics searches against a particular database [34, 35]. This multifaceted approach has been successful, but is labor-intensive, as it requires a separate genomic sequencing step for each sample. The ComPIL and Blazmass methodology uses existing sequence data and can scale to accommodate new data generated by RNASeq and other methods. We feel that a major strength of ComPIL is in identifying unrelated and unexpected peptide sequences that are present in a sample, as we observed with human adenovirus 5 proteins in our HEK293 samples (**Table S2**) or Influenza proteins infecting a human tissue culture cell line.

A previous study directly measuring influenza virus H1N1 infection in Madin-Darby canine kidney cells [36] showed a direct relationship between the infection time of the Calu-3 cells and an increase in Influenza A proteins, in agreement with our results. Furthermore, the relative abundances of the observed influenza proteins generally agree with the previous study, where M1 (matrix protein) and NP (nucleoprotein) are the most abundant viral proteins after >10 hrs post infection and M2 is not observed. Even though the exact Influenza strain that was used to infect the cells was not in the ComPIL database, we were able to detect most Influenza proteins due to high sequence conservation between the Anhui Influenza A strain used to infect human the Calu-3 cells and related strains contained in the database. Importantly, these peptides specifically mapped only to Influenza A proteins and no peptides erroneously matched Influenza B proteins. We re-searched the data using a limited, tailored database consisting of only human and the Anhui Influenza A proteomes and confirmed the above results (**Figure S11**). Importantly, our reanalysis of this publicly available proteomic data is a proof-of-principle demonstration for the employment of a large database proteomic search, where potential organismal contaminants in a biological sample would be identified and would pass undetected with the use of a smaller, focused protein database.

The direct detection of an abundant, conserved, and universally expressed protein produced by the human gut microbiome that contain DUFs highlights a primary utility of untargeted metaproteomics as presented here. We originally focused on proteins containing DUF PF09861, as peptides corresponding to this DUF were ubiquitous among all healthy human stool samples and this domain was unannotated in the Pfam database. Despite our efforts to correlate DUF PF09861 with previously annotated functions, all detected peptides collected from the stool samples only matched proteins with unknown functions. Proteins containing this DUF have poor sequence similarity, but a subset have previously been identified as lactate racemases [37]. While previously known to be primarily found in *Eubacterium* and *Clostridium* genomes, DUF PF09861-containing proteins were under-appreciated with regard to their ubiquitous expression within the context of the healthy human gut microbiota. Such findings support the use of metaproteomics with ComPIL databases as a complementary methodology to metagenomics, as DNA sequencing approaches are incapable of elucidating protein abundance. Our unbiased proteomic searches against large ComPIL databases will continue to shed light on universally and/or differentially expressed microbiome proteins in health and disease and will promote new avenues for human gut microbiome exploration.

The analysis of complex, relatively uncharacterized biological samples such as the human gut microbiome is an important application of our ComPIL database and is the primary driving force for the expansion and improvement of proteomic search databases. To assess the relationship between biological and technical replicates, we performed principal component analysis (PCA) on the samples using spectral counts in protein clusters as features. Notably, PCA reveals excellent grouping of technical replicates for each biological sample (**Figure S12**). The ComPIL database and search methodology can now be employed to quantify microbiome samples across healthy individuals as well as gastrointestinal disease cohorts. Results from these studies will aid in determination of protein family and protein functional conservation among constituents of the microbiota of hosts, and hence, whether conservation in function is preserved in the face of diversity of the composition of the microbiota within the microbiomes of different individuals, populations, and diseases.

We anticipate that ComPIL will allow for large-scale proteomic annotation of systems with extensive protein variation, including human microbiomes. As we transition from the concept of a single ‘reference’ genome or proteome for a species, proteomic methods must adapt to incorporate variant sequences to improve their accuracy and depth of protein identification. We expect that the ComPIL approach will prove useful for future metaproteomic studies in a variety of research areas as publicly-available, high-quality gene and predicted protein sequence data continue to accumulate at an ever-increasing rate. Metaproteomic studies incorporating unbiased database libraries will provide an essential complement to current metagenomic studies and outline new methodologies that are required to construct a foundation on which to map the complexity of the host:microbiome protein interaction network and promote new avenues for human gut microbiome exploration.

### Ethical consideration

All volunteers provided informed consent and the collection and use of human samples were approved by the TSRI Institutional Review Board with reference number IRB-14-6352.

### Availability

Blazmass-ComPIL: https://github.com/sandipchatterjee/blazmass_compil Project home page: https://bitbucket.org/sulab/metaproteomics The mass spectrometry proteomics data have been deposited to the ProteomeXchange Consortium (http://proteomecentral.proteomexchange.org) via the PRIDE partner repository [38] with the dataset identifiers <PXD003896> and <PXD003907>. Programming language: Python, Java, Bash

### Description of Additional Files

Additional file 1: This file contains supplementary figures (PDF 1.4 MB)

## Declarations

### Authors’ contributions

DWW, AIS, JRY conceived of the project. SC designed ComPIL, SKP and JRY designed and wrote Blazmass, and SC, GSS, SKP integrated ComPIL into Blazmass. SC performed wet lab experiments and collected tandem MS data. SC and GSS analyzed tandem MS data. AIS and GSS contributed to peptide mapping and functional analysis. JCD assisted with technical aspects of large database generation, parallelization, and queries. SC, GSS, and DWW wrote the manuscript and all authors contributed to the editing of the manuscript

## Acknowledgments

We thank M. Elsliger, C. Wu, M. Nanis, and W. Shipman for critical suggestions on database generation and performance testing, P. Thuy-Boun for valuable feedback and discussions, and A. Wang, D. Steinmetz, and J. Moresco for technical assistance with sample preparation and mass spectrometry instrumentation.

## Open Access

This article is distributed under the terms of the Creative Commons Attribution 4.0 International License, which permits unrestricted use, distribution, and reproduction in any medium, provided you give appropriate credit to the original author(s) and the source, provide a link to the Creative Commons license, and indicate if changes were made. The Creative Commons Public Domain Dedication waiver applies to the data made available in this article, unless otherwise stated.

## Abbreviations

ComPIL: Comprehensive Protein Identification Library; DUF: domain of unknown function; FDR: false discovery rate; MS: mass spectrometry; MS/MS: tandem mass spectrometry; MudPIT: multidimensional protein identification technology; PSMs: peptide-spectrum matches

